# Accessing Meaning During Encoding Shapes Subsequent Memory Reactivation in the Brain

**DOI:** 10.1101/2025.11.02.686134

**Authors:** Xiaoxi Qi, Marc N Coutanche

## Abstract

Semantic access can occur in many forms, ranging from retrieving specific exemplars to linking concepts with related meanings. We hypothesized that accessing meaning at different levels during encoding would lead to distinct neural reactivation patterns during later memory retrieval. Human participants learned novel word–image associations through prompts that queried item-specific, categorical, or thematic features, while undergoing fMRI. Multivariate pattern analysis shows that encoding histories leave distinct neural signatures even when behavioral performances is equivalent. Encoding history could be decoded by classifiers from recognition activity across the ventral visual stream and higher-order regions. Category-based encoding reduced item-specific reinstatement in perceptual regions, while item-based encoding modified cross-phase item-level similarity in the anterior temporal lobe, suggesting a transition from detailed to integrative conceptual representations. Together, these findings demonstrate that semantic access during encoding shapes how memories are neurally reinstated to integrate perceptual and conceptual information across systems supporting semantic and episodic memory.

---

Memory is not a passive recording of experience. Instead, what we remember and how we remember are fundamentally shaped by how information was processed during encoding. Encoding determines which features are attended to, which associations are activated, and how a stimulus is interpreted within its context (Rissman & Wagner, 2012). This principle is well-established in episodic memory research: attentional focus during encoding enhances memory for attended features while reducing retention of unattended ones (Aly & Turk-Browne, 2016; Chun & Turk-Browne, 2007). Importantly, these encoding effects leave neural traces. Successful retrieval often involves the reinstatement of neural patterns active during encoding, known as pattern reactivation or representational reinstatement (Bruett et al., 2020; Xue, 2018). This similarity between encoding and retrieval neural patterns depends on what information was emphasized and how it was represented neurally (Richter et al., 2016).

Therefore, memory reactivation reflects not the full event, but the subset of features emphasized during encoding. Whether a similar principle applies to semantic memory remains unclear: does the way a concept is accessed during encoding shape how it is later reinstated? If encoding context determines which episodic features are bound into memory, it may also influence which semantic features are re-engaged when a concept is remembered. Here, we investigate this possibility by varying the level of semantic granularity through which a concept is accessed during encoding and examining whether these different forms of semantic processing leave distinct persistent neural traces. Contemporary theories view concepts as distributed representations that integrate perceptual, functional, and relational features across multiple systems (Barsalou, 2008; Ralph et al., 2017a). Although their structure is relatively stable, the aspects of a concept that are activated can shift depending on task and context (Lebois et al., 2015; Yee & Thompson-Schill, 2016). Therefore, conceptual access is dynamic, and one aspect in which it can vary is *semantic granularity*.

Semantic granularity refers to how a concept can contain meanings in different levels of abstraction, each emphasizing distinct features and relational structures. Item-level processing centers on perceptual and functional features that distinguish one instance from another (e.g., *Pluto* the dog). This focus enhances fine discrimination, which is associated with hippocampal pattern separation mechanisms (Xue et al., 2010), and also engages object-specific regions such as the perirhinal cortex (PrC) and anterior temporal lobes (ATL) (Bussey et al., 2006; Lee et al., 2019). Category-level processing organizes concepts based on shared taxonomic features (e.g., dog as an animal), engaging ventral temporal cortex (VT) where domain-selective patterns emerge for conceptual categories (Binder & Desai, 2011; Mahon & Caramazza, 2009). Theme-level processing captures associative or event-based relationships, where meaning emerges from co-occurrence or complementary functional roles (e.g., dog, leash, park). Thematic relations rely on regions such as the ventrolateral prefrontal cortex (vlPFC) supporting schema-based retrieval and event integration (Binder & Desai, 2011; Ranganath & Ritchey, 2012). Previous studies have contrasted these levels of semantic granularity primarily in semantic judgment or comprehension tasks, showing that context determines which type of relation is most accessible (Estes et al., 2011; Lin & Murphy, 2001; Sachs et al., 2008). These three levels engage distinct cognitive processes and neural systems, but whether encoding manipulations targeted at these levels leave lasting representational traces remains unclear.

Here, we manipulated semantic granularity incidentally through question framing during novel word learning. Participants learned new labels for familiar concepts while answering prompts that directed processing toward one of three semantic levels: item-specific features (e.g., Does this bark?), category membership (e.g., Is this an animal?), or thematic relations (e.g., Is this related to parks?). When a new label is paired with a concept, the semantic question effectively “binds” the label to the conceptual features emphasized at the targeted level, creating distinct encoding-history traces across conditions. These encoding effects were expected to arise naturally from the semantic processing demands induced by each question type, consistent with evidence that individuals can rapidly integrate new information into memory structures through incidental semantic inference (Coutanche & Thompson-Schill, 2014; Zaiser et al., 2022). We assessed how these histories persisted by training classifiers and quantifying neural representational strength during final recognition. Neural representational strength was quantified using *Encoding-Recognition Similarity* (ERS), which captures the reinstatement of encoding-phase representations during subsequent recognition. We predicted that representational strength should be greatest when encoding history aligns with the semantic structure being tested (e.g., item encoding enhances item-level patterns). These effects were expected to be associated with functionally relevant systems (e.g., item-related patterns in ATL, category-related patterns in VT, and theme-related patterns in vlPFC).

## Results

### Behavioral Accuracy

Participants successfully learned the word-image pairings, showing recognition accuracy well above chance (M = 0.69, SD = 0.21, chance= 0.25, t(29) = 11.47, p<0.001). To examine the effects of encoding history on memory performance, we conducted a rmANOVA on recognition accuracy. There was no significant effect of encoding history (F(2,58) = 1.00, p = 0.37), suggesting that neural differences are not resulting from systematic unequal memory performance across encoding histories. Mean performance did not differ significantly across item-, category-, and theme-based encoding histories (item: M = 0.71, SD = 0.19; category: M = 0.70, SD = 0.23; theme: M = 0.67, SD = 0.26), indicating comparable overall behavioral learning across encoding history conditions.

### Classifying Encoding History from Recognition Patterns

#### Whole-brain Searchlight Classification

To identify where neural activity patterns during recognition carried information about encoding history, we conducted a whole-brain searchlight classification analysis (chance = 33%).

Five significant clusters exceeded the cluster-corrected threshold, revealing regions where local patterns successfully distinguished between encoding histories. These clusters were located bilaterally in EVC and extended anteriorly along the ventral visual stream, including the fusiform gyrus, posterior occipital regions, and a smaller cluster overlapping with the angular gyrus (Fig. 1; Table 1).

**Table 1.** Significant Searchlight Clusters that Decoded Encoding History Based on Recognition Patterns.

| Cluster # | Regions | Cluster Size<br>(voxels) | X | Y | Z |
| --- | --- | --- | --- | --- | --- |
| 1 | Right lingual gyrus | 2332 | 16 | -77 | -5 |
| 2 | Right superior occipital gyrus | 442 | 22 | -72 | 34 |
| 3 | left angular gyrus | 411 | -42 | -67 | 34 |
| 4 | left middle occipital gyrus | 353 | -37 | -77 | 3 |
| 5 | left fusiform gyrus | 300 | -34 | -80 | -16 |

*Notes.* Cluster sizes reflect the number of voxels surviving correction (voxel-wise $p < .005$ , cluster-level $\alpha = .05$ ). Center of mass coordinates are reported in MNI space (LPI orientation).

**Figure 1.**
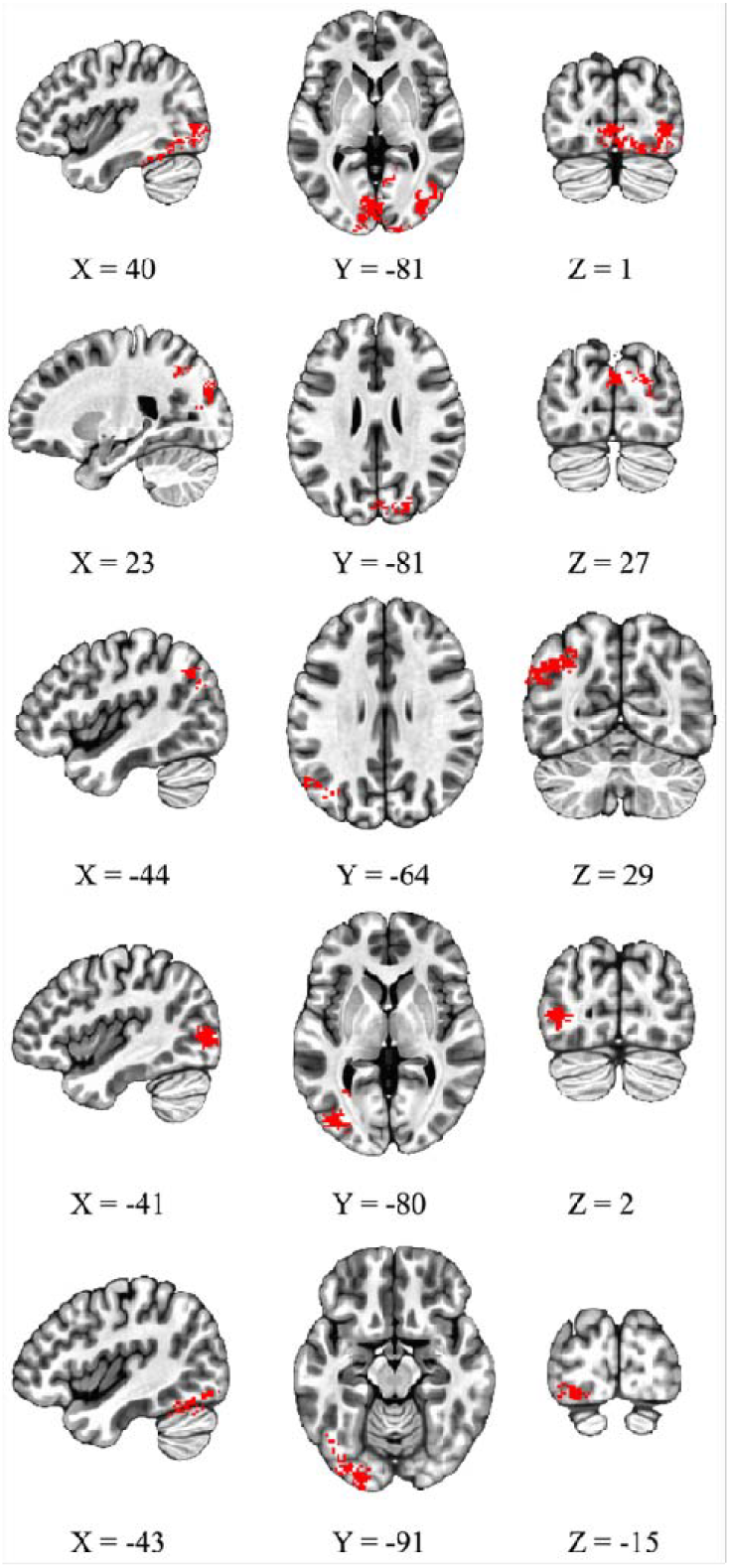
Significant Clusters from Whole-brain Searchlight Classification Analysis. *Notes*. Example slices showing significant clusters overlaid on a MNI template brain in LPI coordinate space. Each row corresponds to one cluster, presented in the same order as listed in Table 2. The displayed coordinates were manually selected to optimize the visibility of each cluster.

**Figure 2.**
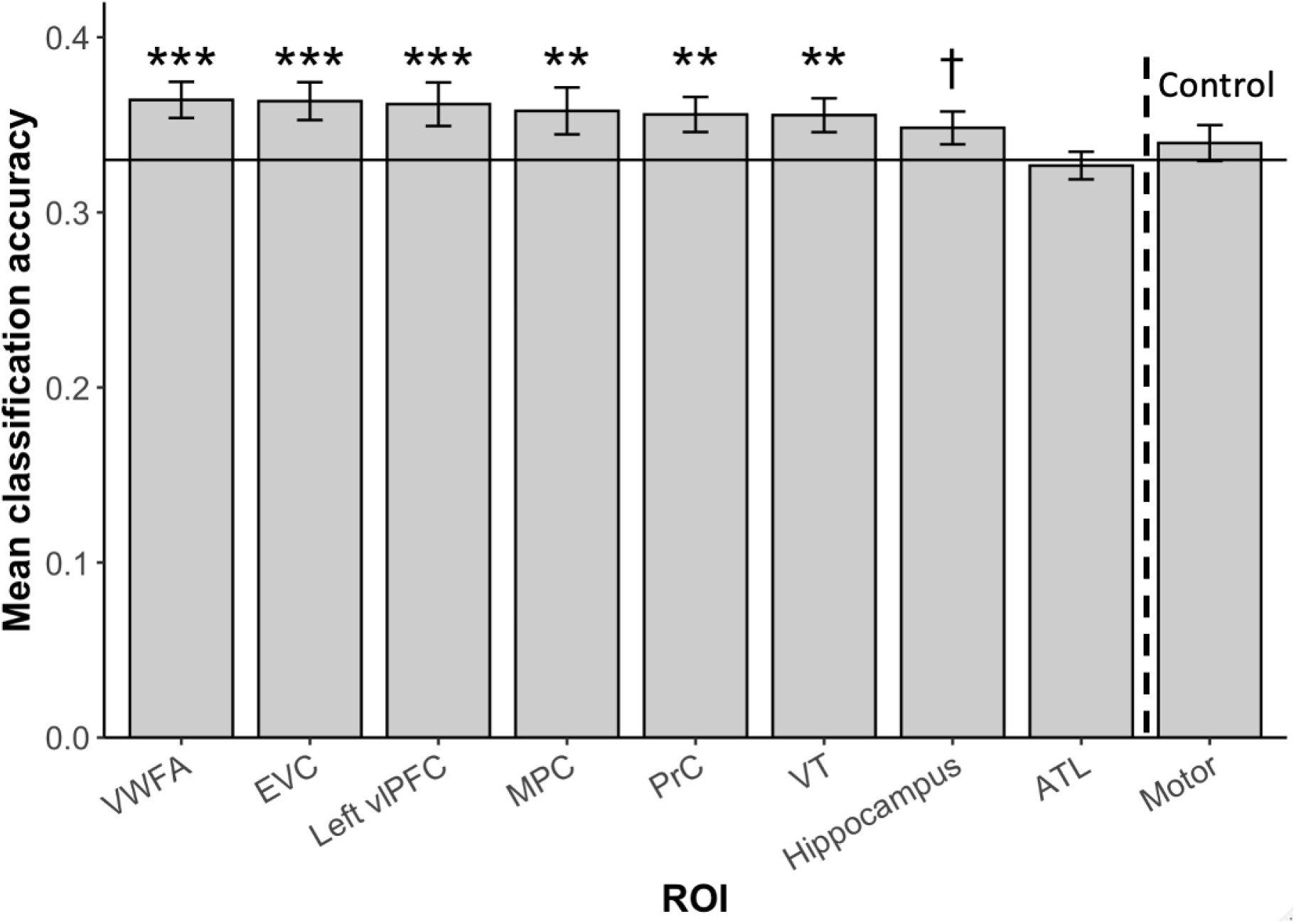
Three-way Classification of Encoding History in All Recognition Data. *Notes*. The classifiers were trained and tested on all trials to decode prior encoding histories (item vs. category vs. theme). Error bars represented standard errors. The horizontal line represents the chance level of 33%. The dashed vertical line is used to separate the ROIs and the control region. † p < .1, *p<.05, **p<.01, ***p<.001.

#### ROI-Based Classification

Within predefined ROIs, Gaussian Naive Bayes (GNB) classifiers achieved above-chance accuracy in multiple regions, confirming that encoding history could be decoded from recognition activity patterns. After FDR correction, significant decoding was observed in VT (M = 35.55%, SE = 0.97%, p = 0.001, p_adj_= 0.002), PrC (M = 35.59%, SE = 1.00%, p = 0.001, p_adj_ = 0.002), MPC (M = 35.80%, SE = 1.34%, p = 0.002, p_adj_ = 0.003), left vlPFC (M = 36.18%, SE = 1.25%, p < 0.001 p_adj_ < 0.001), EVC (M = 36.35%, SE = 1.08%, p < 0.001, p_adj_ < 0.001), and VWFA (M = 36.42%, SE = 1.03%, p < 0.001, p_adj_ < 0.001). The hippocampus showed a marginal effect that did not survive correction (M = 34.83%, SE = 0.94%, p = 0.042, p_adj_ = 0.054). ATL (M = 32.67%, SE = 0.79%, p = 0.79, p_adj_ = 0.79) and the motor cortex as a control region (M = 33.96%, SE = 1.02%, p = 0.24, p_adj_ = 0.27) did not show above-chance classification.

### Encoding–Recognition Similarity (ERS)

We examined whether encoding history influenced the reinstatement of semantic representations during recognition. A rmANOVA on ERS revealed significant interactions between encoding history and semantic representation structure in ATL (F(4, 116) = 2.76, p = 0.031) and EVC (F(4, 116) = 8.00, p < 0.001) (Figure 3). In ATL, item-specific reactivation was lower following item-based encoding than after theme-based encoding (t(29) = –2.63, p = 0.014, p_adj_ = 0.041). In EVC, category-based encoding produced weaker item-specific reactivation than both item-based encoding (t(29) = –3.06, p = 0.005, p_adj_ = 0.014) and theme-based encoding (t(29) = –2.68, p = 0.012, p_adj_ = 0.018), suggesting that emphasizing shared categorical features may hinder the reinstatement of item-specific patterns.

**Figure 3.**
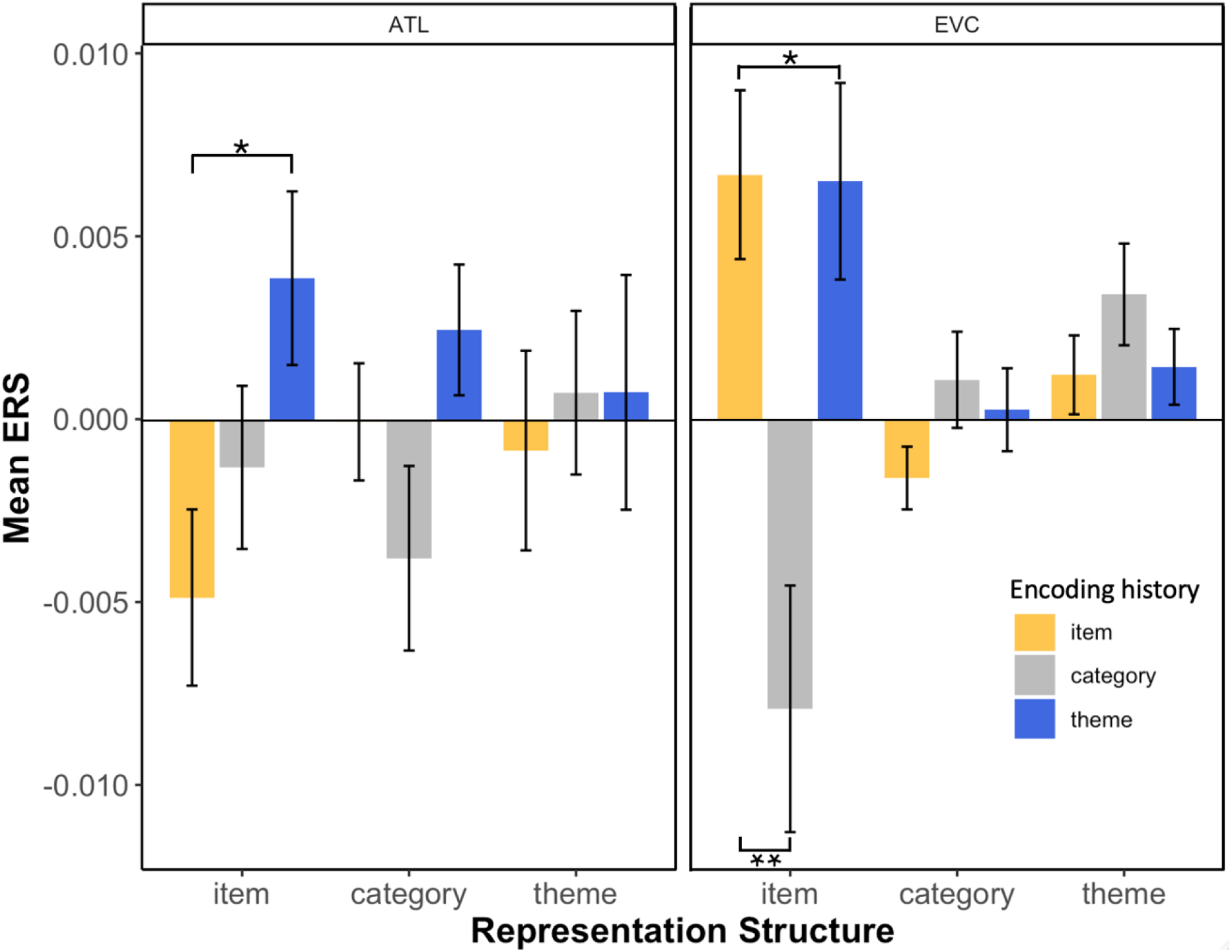
Significant Encoding History x Representation Structure Interaction on ERS. *Notes*. For all stimuli, interactions between representation strucutre and encoding history were found in ATL and EVC. Pairwise comparison results are FDR corrected. *p < .05, **p < .01.

Several additional regions showed main effects independent of the interaction. Left vlPFC (F(2, 58) = 4.63, p = 0.014), VT (F(2, 58) = 4.38, p = 0.017), and hippocampus (F(2, 58) = 3.63, p = 0.033) showed main effects of encoding history, indicating overall differences in reactivation strength across encoding histories. A main effect of representation structure was found in PrC (F(2, 58) = 3.83, p = 0.027) and EVC (F(2, 58) = 5.98, p = 0.004), showing that reactivation strength varied depending on semantic structure. These effects were not followed up with post hoc comparisons, as they were not central to our hypotheses and were qualified by interactions in key ROIs. However, they offer additional evidence that both encoding history and semantic representation structure can independently influence the strength of neural reactivation during recognition.

## Discussion

This study investigated how accessing different semantic aspects of a concept while encoding a novel word shapes the strength and structure of its subsequent neural reactivation. By manipulating semantic granularity through question framing, we created distinct encoding histories that bound each novel word to one of three semantic levels: item, category, and theme. Participants successfully learned these pairings, and recognition accuracy did not differ across encoding histories, confirming that observed neural effects reflect differences in semantic processing rather than general memory strength. Across multiple analyses, encoding histories left measurable and distributed traces on neural activity during recognition, demonstrating that the semantic information emphasized during encoding leaves a lasting influence on how memories are represented and reactivated.

Classification analyses showed that encoding history could be decoded from distributed activity during recognition. Whole-brain searchlight classification identified ventral visual clusters extending from EVC through the fusiform gyrus, consistent with the ventral stream’s progression from perceptual to conceptual processing (Binder & Desai, 2011; Grill-Spector & Weiner, 2014). ROI analyses further confirmed above-chance decoding performance in EVC, VWFA, VT, MPC, PrC, and left vlPFC, with marginal effects in the hippocampus. The distributed pattern supports models proposing that conceptual retrieval depends on a broad cortical network integrating sensory, semantic, and control systems (Binder et al., 2009; Ranganath & Ritchey, 2012). In contrast, ATL, as a key semantic hub, did not show reliable decoding. This lack of effect likely reflects its representational structure rather than reduced sensitivity to encoding history. ATL codes semantic information in graded and overlapping patterns that are difficult to separate linearly (Ralph et al., 2017a). As a result, our classifier may underestimate, while similarity-based analyses may be more capable of revealing ATL’s contribution (Diedrichsen & Kriegeskorte, 2017; Kriegeskorte et al., 2008). Indeed, ATL showed significant effects in representational similarity analysis (discussed below), indicating that encoding history altered the representational structure rather than how separable its representations were. These findings demonstrate that semantic information engaged during encoding leaves persistent traces throughout the perceptual-conceptual hierarchy, modifying both the distribution and internal structure of recognition activity patterns. This provides neural evidence that the semantic processing context becomes integrated into subsequent memory representations, just like episodic ones.

ERS, operationalized as cross-phase similarity between encoding and recognition, revealed significant interactions between encoding history and representational structure in EVC and ATL. In EVC, category-based encoding reduced item-specific reactivation compared to item- and theme-based encoding, suggesting that emphasizing shared categorical features suppresses detailed perceptual reinstatement. This pattern is consistent with EVC’s role in the reinstatement of perceptual details during episodic retrieval, particularly for visually rich stimuli (Danker & Anderson, 2010). In ATL, item-based encoding produced negative ERS values significantly lower than theme-based encoding, indicating that identical items were represented differently across phases. This likely reflects ATL’s role in integrating modality-specific inputs into coherent conceptual representations (Chiou & Lambon Ralph, 2016). When encoding emphasizes fine perceptual detail (item-based encoding), ATL activation may vary more across contexts; however, ATL activity converges onto an abstract and amodal representation during retrieval (Chiou & Lambon Ralph, 2016; Peelen & Caramazza, 2012; Rice et al., 2015). On the other hand, theme-based encoding, which already highlights broad associative information, may produce greater cross-phase overlap in this integrative semantic hub region.

In summary, the present findings show that the semantic granularity engaged during encoding (item, category, theme) shapes how memories are later represented in the brain, by influencing the reactivation of memory representations. Distinct encoding histories left measurable traces across perceptual and conceptual systems, with perceptual regions such as EVC showing reduced item-specific reinstatement after category-based encoding and integrative hub regions like ATL showing shifts in representational patterns across phases, especially after item-based encoding. Together, these findings provide new insights into how the level of semantic granularity shapes memory reactivation, informing broader theories of memory representation, conceptual knowledge, and neural mechanisms supporting memory retrieval.

## Methods

### Participants

Based on a power analysis using effect sizes from a pilot study, a minimum of 23 participants is needed to achieve a power of 0.8. Thirty-six participants were recruited for the study. We employed strict thresholds for behavioral accuracy in order to ensure participants learned the introduced information (>55% for localizer or >75% for re-exposure). This removed 5 for low behavioral accuracy on behavioral tasks, and one for an incomplete scan due to claustrophobia. Therefore, 30 participants (22 females; age: Mean (M)= 24.1, standard deviation (SD)= 6.4) were included for data analyses. Participants provided written informed consent and were compensated for their time. All participants, aged between 18 and 39 years old, were right-handed with normal or corrected-to-normal vision, and no report of psychiatric or neurological conditions. Participants met the standard MRI safety requirements of the neuroimaging center, including an absence of metal in the body and a low likelihood of pregnancy. All participants were native English speakers with no previous experience with German and/or Dutch languages. This study was approved by the university’s Institutional Review Board.

### Stimuli

Participants learned associative pairings of object images with novel words. Novel words were selected Dutch words, each was 3 to 8 letters long, based on a normalized dataset (Tokowicz et al., 2002). All selected words were concrete nouns with single English translations and exhibited low form similarity with their English counterparts, according to ratings from 24 Dutch-English bilinguals. The study included 24 images representing six themes (Vet, Picnic, Ocean, Garbage, Winter, and Summer) and four categories (Animals, Food, Furniture, and Tools). Each image was presented four times across two runs (2 trial each run) during encoding, restudy, and recognition tasks. Images were scaled to the same size (250 × 250 pixels) and placed on a white background.

### Experimental procedure

After providing consent, participants completed out-of-scanner behavioral category training. Five tasks were administered during the fMRI scan (Taks sequence is presented in Figure 4). The trial order for in-scanner tasks was pseudorandomized in advance, following the optimal sequence determined by the jitter optimization program Optseq 2 (https://surfer.nmr.mgh.harvard.edu/optseq/). After the scan, participants completed an out-of-scanner cued recall task.

**Figure 4.**
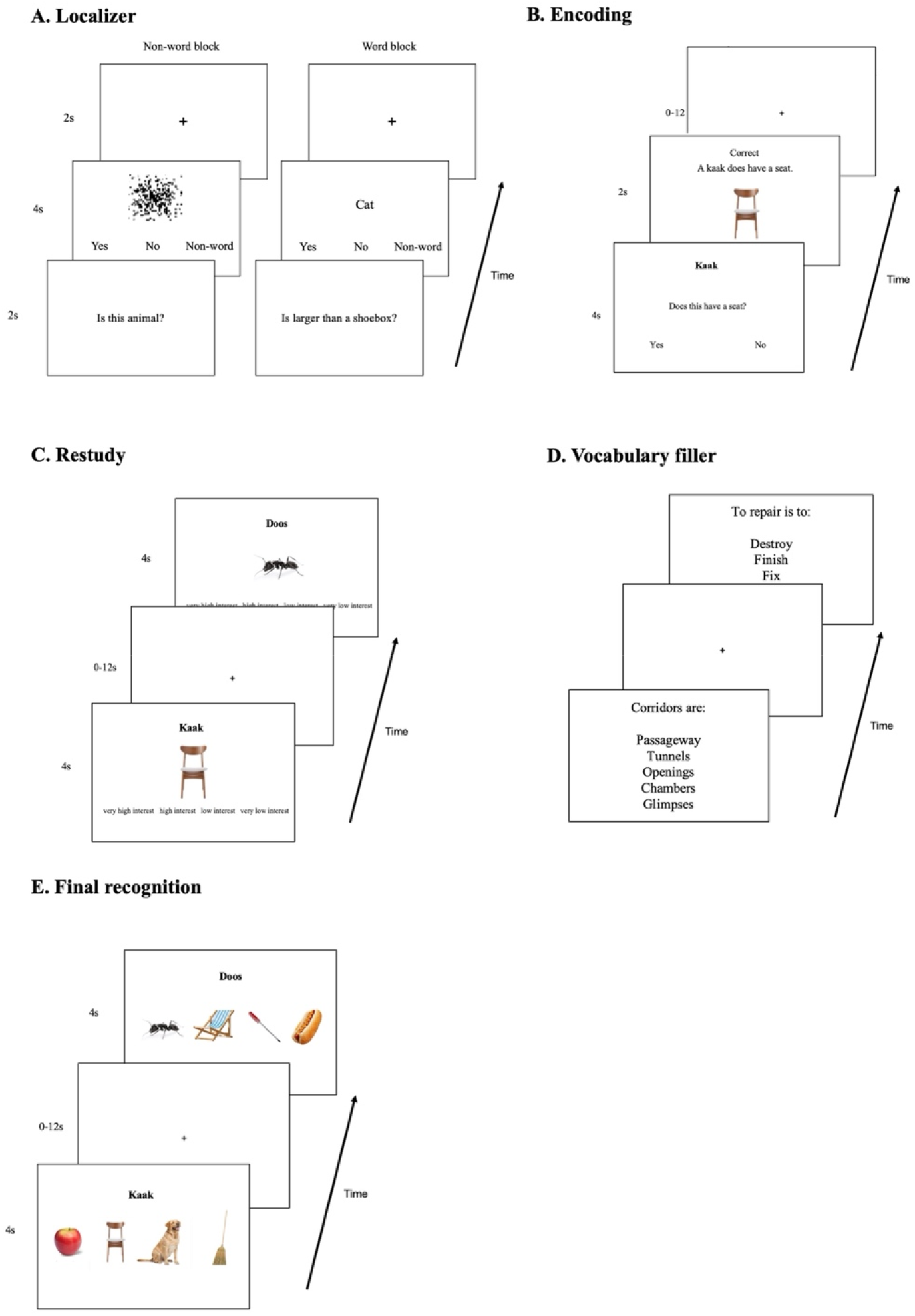
In-scanner task sequence. *Notes*. (A) Localizer task included alternating word and non-word blocks. Participants responded to a question presented at the beginning of each block (e.g., “Is this an animal?”), followed by real words or scrambled images. (B) During encoding, participants answered a question targeting one of three semantic levels (e.g., “Does this have a seat?”) about each item to learn its paired novel name. Corrective feedback followed each response. (C) During restudy, participants viewed encoded word–image pairs and rated their interest. (D) The vocabulary filler task, shown during the anatomical scan, was selecting a synonym for a given word from 4 options. (E) During final recognition, participants matched each novel word to one of 4 images, which included the correct paired image and three other learned images from different categories and themes.

#### Category training

We familiarized participants with the definitions of the object categories used in the study: animal, food, furniture, and tools. Twelve known English words (3 for each category) were presented as examples, followed by a question targeting the category level of each word (e.g., “Is this food?”). Participants received feedback after each question (“Onion is a food. Foods can be consumed”). Words presented in this task did not overlap with items later paired with novel names.

#### Localizer

We identified the visual word form area (VWFA) for each participant with a 10-minute localizer across two runs, each containing six blocks. There were two block types: word and non-word. At the start of each block, participants saw a yes/no question targeting a specific semantic level (e.g., “Is this an animal?”). In word blocks, participants viewed a series of real words and responded “yes” or “no” based on the question shown at the beginning of the block. In non-word blocks, participants viewed scrambled images and pressed a third button to indicate a “non-word” response. Each block began with the question screen presented for 2 seconds, followed by eight stimuli shown for 4 seconds each. Stimuli were separated by two seconds of fixation. This task was used to identify regions sensitive to semantic granularity and word processing. To avoid interference, the words and questions used in this localizer task did not overlap with the stimuli used in the main tasks.

#### Encoding

Participants were asked to learn 24 pairs of item images and their novel names (Dutch words) by answering yes/no questions about the item, targeting one of three semantic levels. Each pair and testing question was presented for 4 seconds, twice per run (4 times across 2 runs), with the correct answer to the testing question being “yes” once and “no” once. Testing questions invoked the target level of semantic granularity: item questions (e.g., “Is *kaak* bigger than a shoebox?”) probed stimulus-specific details, category questions (e.g., “Is *kaak* furniture?”) probed the semantic category to which it belongs, and theme questions (e.g., “Is *kaak* related to a classroom?”) probed its thematic relationships. We provided feedback after responses for 2 seconds (e.g., “Correct. A *kaak* is a chair.” / “Incorrect. A *kaak* is not an animal.”). Past findings on the testing effect suggested that corrective feedback often enhances subsequent memory performance (Kang et al., 2007). Pairing randomization was counterbalanced across participants. If the participant did not respond within 4 seconds, the trial was considered “incorrect.” After the feedback, there was a jittered interstimulus interval of 0-12 seconds (average of 2 seconds).

#### Restudy

Participants were presented with word-image pairs they encoded earlier and informed that they need to memorize the presented information. Each pair was presented for 4 seconds, 4 times across two runs (twice per run). To maintain participants’ engagement with the task, they rated their interest in each pair on a four-point scale using an MRI-compatible button box. There was a jittered interstimulus interval of 0-12 seconds (average of 2 seconds).

#### Vocabulary Filler

This task was administered during the anatomical scan. Participants were presented with a target word and asked to select which of 4 words most closely matched the target in meaning. This task was not relevant to the main analyses and was used to prevent additional rehearsal associated with the learned stimuli.

#### Recognition

Participants were asked to recognize the correct image associated with each novel word from 4 previously encoded images. The 4 images included all 4 object categories and 4 different themes. Each novel word appeared twice in each run, with each trial lasting for 4 seconds. There was a jittered interstimulus interval of 0-12 seconds (average of 2 seconds).

#### Cued Recall

After the scan, participants completed an out-of-scanner final cued recall task, typing the associated Dutch word while viewing each previously paired image. Participants were instructed to provide their best guess when spelling the words. Each trial ended when participants indicated they had finished typing by pressing the “enter” key. This task was not analyzed here.

### Image acquisition

All scanning was performed on a Siemens 3-T Prisma MRI scanner equipped with a mirror device to perform fMRI stimuli presentation. Anatomical T1-weighted whole-brain MRI images were collected between re-exposure and final recognition phase (TR =1.540 s, TE =3.04s, voxel size = 1.0 × 1.0 × 1.0 mm). Data collection for all tasks took over eight functional runs in total. All functional runs had whole-brain coverage with 2.0 × 2.0 × 2.0 mm voxel size and a TR of 2 seconds. A predetermined jitter between 0 and 12 seconds (mean = 2 seconds) was used in tasks after the localizer to extract the signal from the rapid event-related design, with the optimal sequence determined using Optseq 2 (https://surfer.nmr.mgh.harvard.edu/optseq/).

### Image preprocessing

Imaging data were processed using the Analysis of Functional NeuroImages (AFNI) software package (Cox, 1996). Structural scans were skull-stripped and standardized to MNI standard space. Motion correction was applied to all functional images, which were registered to a mean functional volume. One participant was removed due to excessive motion. To conduct multivariate analyses, beta coefficients were calculated for each trial using the Least Squares-Separate (LSS) method (Mumford et al., 2014). Each voxel’s beta coefficients were z-scored across the time course of each run.

### Regions of interest

We defined eight regions of interest (ROIs) based on semantic processing and word learning literature (Figure 5). Bilateral VT cortex is involved in encoding complex visual stimuli and includes category-selective regions (Haxby et al., 2011). To define VT, we used anatomical criteria based on prior literature: extending from 70 to 20 mm posterior to the anterior commissure in Talairach coordinates, incorporating the lingual, fusiform, parahippocampal, and inferior temporal gyri (Haxby et al., 2001). The ATL acts as a hub for semantic memory by integrating information across multiple modalities (Lambon Ralph, 2014; Rice et al., 2015). We defined ATL using coordinates from prior studies, forming 6 mm-radius spheres at MNI coordinates (after conversion): 43, 13, −25; −42, 13, 23 (Coutanche & Thompson-Schill, 2015). The left vlPFC is active during speech comprehension and semantic retrieval (Nozari & Novick, 2017), and it was defined using Brodmann areas BA44, BA45, and BA47 from a Brodmann atlas (Mai, 2017).

**Figure 5.**
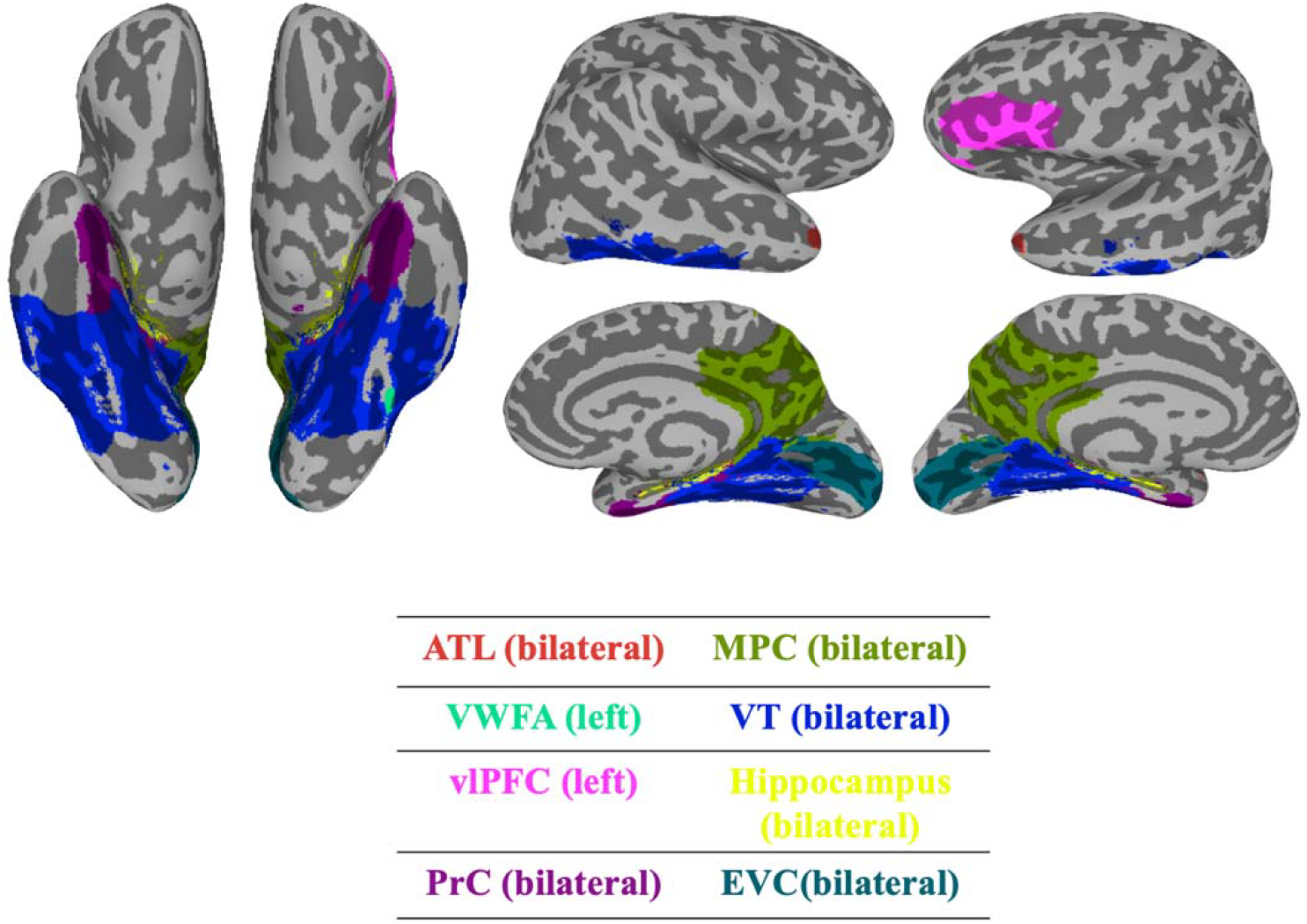
ROIs on a MNI template brain. *Notes*. Surface visualization of all regions of interest (ROIs) used in the study, shown on a standard MNI brain. ROIs include bilateral ATL, VT, EVC, MPC, PrC, hippocampus, and left-lateralized VWFA and vlPFC. Each ROI is color-coded and labeled below for reference.

Early visual cortex (EVC) has been modulated by top-down semantic and mnemonic processes (e.g., Ehrlich et al., 2024; Sergent et al., 2011) and we have previously found that EVC patterns can be shifted by learning perceptually-relevant dimensions (such as real-world size; Coutanche & Thompson-Schill, 2019). EVC was defined using Brodmann areas BA17.

The VWFA demonstrates sensitivity to orthographic regularities, responding to both real words and pseudowords (Chen et al., 2019; Cohen et al., 2002). We identified regions sensitive to the word vs. non-word contrast using the localizer task, near the left fusiform gyrus. Then we created a 6 mm-radius sphere around the coordinate of the peak activity.

Additionally, the hippocampus and PrC are crucial for memory processes, with encoding activation in PrC predicting later item recognition (Davachi et al., 2003). The medial parietal cortex (MPC) was included due to its association with successful memory recall and its activation in response to visual stimuli from various categories, particularly those involving people and places. The functional organization of the MPC also shows connectivity to the VT cortex (Silson et al., 2019). These three regions were defined using Freesurfer anatomical atlas parcellations. The MPC specifically includes the precuneus, subparietal sulcus, and posterior cingulate gyrus.

### Behavioral analyses

To assess whether recognition performance was influenced by encoding history, we conducted a repeated measures ANOVA with encoding history (Item, Category, Theme) as within-subject factor. Recognition accuracy was calculated as the proportion of correctly identified image-word pairs across conditions.

### Mapping Neural Representations of Semantic Granularity

To identify brain regions sensitive to intrinsic semantic structure (item, category, theme), we compared neural representational similarity matrices (RSMs) during recognition with model RSMs capturing each semantic level. Correlations were computed using second-order Spearman’s ρ and evaluated with permutation tests per ROI. A whole-brain searchlight identified additional regions showing reliable correspondence to model structures (voxelwise p < .005, cluster-wise α = .05, corrected with AFNI ClustSim) (See details and results in SI Appendix).

### Classifying Encoding History from Recognition Patterns

To determine whether recognition-phase activity patterns carried information about prior encoding history, we performed multivariate pattern classification using a Gaussian Naive Bayes (GNB) classifier. The classifier was trained to distinguish between the three types of encoding history (item, category, theme; chance = 33%) using a 2-fold cross-validation procedure, in which each recognition run served as training and testing data across folds. Analyses were performed both within predefined ROIs and across the whole brain using a searchlight approach.

Whole-brain analysis was conducted by applying a 6-mm radius spherical searchlight to each voxel, and classification accuracy was assigned to the center voxel, producing participant-level accuracy maps. These maps were normalized to MNI space and entered into a group-level one-sample t-test to test whether voxel-wise accuracy exceeded chance. Cluster correction was performed using AFNI’s 3dttest++ with ClustSim, with a voxel-wise threshold of p < .005 and a cluster-wise α = .05, corresponding to a minimum cluster size of 267 voxels.

ROI-based classification was conducted within predefined ROIs. Statistical significance was assessed using a nonparametric permutation test, in which class labels were randomly shuffled 1,000 times to generate a null distribution of accuracies. The p-value was computed as the proportion of permutations yielding accuracy equal to or greater than the observed value, and then corrected for False Discovery Rate (FDR).

To make sure that memory performance differences did not drive decoding accuracy, we repeated the analysis using only correctly recognized stimuli (defined as items correctly identified in more than 2 of 4 recognition trials). To match class sizes across encoding histories, we balanced trial counts by subsampling to the smallest available class per participant. Participants with fewer than four correct trials in any encoding history were excluded, resulting in a final sample of 25 participants for this memory-accuracy-controlled analysis.

### Encoding—Recognition Similarity (ERS)

To test whether encoding history influenced the reinstatement of semantic information during later retrieval, we quantified neural pattern reactivation by computing encoding— recognition similarity (ERS). For each phase, beta maps from the two trials of each stimulus were averaged within each run. Then, ERS was computed by correlating each stimulus’s voxel-wise activity pattern from one run during encoding with each run during recognition, resulting in four correlations per stimulus pair (2 × 2 combinations). These correlations were Fisher-Z-transformed and averaged to obtain a single ERS value per stimulus pair, computed separately for each participant and ROI.

To quantify ERS for each semantic representation structure (item, category, theme), we extracted structure-specific averages from the full cross-phase RSMs:

- Item structure: average correlation between identical items across phases (within, e.g., chair_encoding_ – chair_recognition_) compared to average correlation between different items (between, chair_encoding_ – dog_recognition_).
- Category structure: average correlation between items from the same category (within, e.g., chair_encoding_ – table_recognition_) compared with average correlations between items from different categories (between, e.g., chair_encoding_ – apple_recognition_).
- Theme structure: average correlation between items from the same theme (within, e.g., leash_encoding_ – dog treat_recognition_) compared to average correlation between items from different themes (between, e.g., leash_encoding_ – chair_recognition_).

ERS for each representational structure was operationalized as the difference between within-structure and between-structure similarity, producing item-, category-, and theme-structure reactivation scores. These scores were used in a 3 (Encoding History: item, category, theme) × 3 (Semantic Representation Structure: item, category, theme) repeated-measures ANOVA. Post-hoc pairwise comparisons were conducted after significant interaction effects, and p-values were FDR-corrected for multiple comparisons.

## Data availability

The fMRI and behavioral data supporting the findings of this study are available upon request. Source data underlying all figures and statistical results will be made publicly available at the time of publication.

## Code availability

All custom analysis scripts used for data preprocessing, model fitting, and statistical analyses will be released on a public GitHub repository (repository links will be provided upon publication). The code will be made available for academic research use.

